# CFP-1 interacts with HDAC1/2 complexes in *C. elegans* development

**DOI:** 10.1101/451237

**Authors:** Bharat Pokhrel, Yannic Chen, Jonathan Joseph Biro

## Abstract

CFP-1 (CXXC finger binding protein 1) is an evolutionarily conserved protein that binds to non-methylated CpG-rich promoters in humans and *C. elegans*. This conserved epigenetic regulator is a part of the COMPASS complex that contains the H3K4me3 methyltransferase SET1 in mammals and SET-2 in *C. elegans*. Previous studies have indicated the importance of *cfp-1* in embryonic stem cell differentiation and cell fate specification. However, neither the function nor the mechanism of action of *cfp-1* is well understood at the organismal level. To further investigate the function of CFP-1, we have characterised *C. elegans* COMPASS mutants *cfp-1(tm6369)* and *set-2(bn129)*. We found that both *cfp-1* and *set-2* play an important role in the regulation of fertility and development of the organism. Furthermore, we found that both *cfp-1* and *set-2* are required for H3K4 trimethylation and play a repressive role in the expression of heat shock and salt-inducible genes. Interestingly, we found that *cfp-1* but not *set-2* genetically interacts with Histone Deacetylase (HDAC1/2) complexes to regulate fertility, suggesting a function of CFP-1 outside of the COMPASS complex. Additionally we found that *cfp-1* and *set-2* acts on a separate pathways to regulate fertility and development of *C. elegans*. Our results suggest that CFP-1 genetically interacts with HDAC1/2 complexes to regulate fertility, independent of its function within COMPASS complex. We propose that CFP-1 could cooperate with COMPASS complex and/or HDAC1/2 in a context dependent manner.

## Introduction

Chromatin regulation shapes gene activity, which underlies many biological processes including development. Histone modifications are a major form of chromatin modification that play a central role in controlling gene expression (Tessarz and Kouzarides, 2014). The perturbation of these modifications have been associated with developmental defects and diseases including cancer (Shilatifard, 2012, Allis and Jenuwein, 2016, Celano et al., 2018). However, the mechanism by which histone modifications contribute to these events is yet to be explored.

The interplay between the highly dynamic histone modifications can determine chromatin regulation and gene function (Zhang et al., 2015). At enhancer and promoter regions histones are subjected to high turn-over of acetylation or methylation modifications which results in either activation or repression of gene expression (Berger, 2007, Kouzarides, 2007, Thakur et al., 2003). Acetylation of histones by conserved histone acetyltransferases (HATs) such as Gcn5, p300/CBP, sRC/p160 and MYST is related to gene expression. Whereas deacetylation of histone by evolutionarily conserved histone deacetylases (HDACs) is often associated with gene repression (Kouzarides, 2007, Wang et al., 2017, Lau et al., 2016). HDACs form multiprotein complex such as SIN3, NuRD, and CoREST complexes to regulate gene expression (Kouzarides, 2007).

One of the most studied chromatin modifications is histone 3 lysine 4 trimethylation (H3K4me3). H3K4me3 is found at 5’ sites of active genes and is often regarded as an active promoter mark (Barski et al., 2007, Pokholok et al., 2005). Previous studies have shown that the level of H3K4me3 is strongly correlated with gene expression of a subset of genes. These studies have suggested that H3K4me3 could contribute to gene expression by acting as a binding site for chromatin modifiers and transcriptional machinery to facilitate the transcription process (Pena et al., 2008, Wysocka et al., 2006, Ardehali et al., 2011, Howe et al., 2017). Contrary to the role of H3K4me3 in gene expression, growing evidence have suggested that H3K4me3 could play a repressive role in gene expression (Margaritis et al., 2012, Lorenz et al., 2012, Robert et al., 2014). All these findings generated from different organisms suggest that H3K4me3 could play a role in both gene expression and repression in a context dependent manner. Nevertheless, how H3K4me3 contributes to gene expression and repression needs to be explored further.

H3K4me3 is deposited by a Complex Proteins Associated with Set1 (COMPASS) complex (Shilatifard, 2012). The COMPASS complex is evolutionarily conserved from yeast to mammals. In yeast, there is only one complex which is responsible for all forms of H3K4 methylation (H3K4me1, H3K4me2, and H3K4me3), whereas in humans there are six COMPASS complexes: SET1A, SET1B, Mixed Lineage Leukemia (MLL) 1, 2, 3 and 4. SET1A and SET1B are responsible for the majority of H3K4me3 mark deposition, MLL 1 and 2 are responsible for H3K4me3 deposition in a subset of genes, and MLL 3 and 4 are responsible for H3K4me1 (Shilatifard, 2012). In *C. elegans* there are two COMPASS complexes: SET-2/COMPASS, a direct descendent of yeast Set1, and SET-16/COMPASS which is MLL 3/4 ortholog (Li and Kelly, 2011, Simonet et al., 2007). Although Set1 is the key subunit of COMPASS, its associated subunits are also important for assembly and regulation of H3K4 methylation (Krogan et al., 2002, Shilatifard, 2012).

One of the major subunits of the COMPASS complex is CFP1 which is essential for H3K4me3 modifications (Lee and Skalnik, 2005, Skalnik, 2010, Clouaire et al., 2014). CFP1 binds to unmethylated CpG-rich DNA sequences known as CpG islands (CGIs) and helps the recruitment of SET1/COMPASS complex at the promoter region of active genes (Chun et al., 2014, Chen et al., 2014, Thomson et al., 2010, Clouaire et al., 2012, Tate et al., 2010). Previous studies have reported that CFP1 plays an important role in cell fate specification and cell differentiation (Clouaire et al., 2014, Mahadevan and Skalnik, 2016, Skalnik, 2010). However, the exact mechanism by which CFP1 contributes to gene regulation and development is not clear. To understand the role of *cfp1* in gene regulation and development, we have used *cfp-1(tm6369)* and *set-2(bn129) C. elegans* mutants. We discovered that deletion of *cfp-1* or *set-2* results in drastic reduction of H3K4me3 levels and stronger expression of heat shock and salt-inducible genes. Surprisingly, we found that despite both genes being essential for H3K4me3 deposition and gene induction, only *cfp-1* but not *set-2* genetically interacts with HDAC1/2 in *C. elegans* development. This study suggests that in addition to the canonical function of CFP-1 in the H3K4me3 deposition, CFP-1 also cooperates with HDAC 1/2 complexes during *C. elegans* development.

## Results

### CFP-1 is required for fertility and normal growth rate

In yeast and mammals, COMPASS/Set1 is responsible for the majority of H3K4me3. Loss of *SET1* or *CFP1* results in drastic reduction of H3K4me3 levels at 5’ sites of active genes (Lee and Skalnik, 2005, Skalnik, 2010, Miller et al., 2001, Lee et al., 2007, Brown et al., 2017). Similar to mammals and yeast, function of *set-2* (homolog of *SET1)* and *cfp-1* (homolog of *CFP1)* in H3K4me3 deposition is also conserved in *C. elegans* (Simonet et al., 2007, Li and Kelly, 2011, Xiao et al., 2011). To further investigate the role of *cfp-1* in development, *cfp-1(tm6369)* mutants were used in this study. *cfp-1(tm6369)* is a deletion allele, which has 254bp deletion on exon 5 of F52B11.1a.1 (Figure. 1A). This deletion results in a frameshift mutation after 371 aa and a premature stop three codons later. Exon 5 is conserved in all transcripts of the *cfp-1* gene, therefore deletion on exon 5 region of F52B11.1a.1 results in truncation in all the transcripts of *cfp-1* gene (Figure 1A). To confirm that *cfp-1(tm6369)* is a loss of function allele we measured the global level of H3K4me3 in both *cfp-1(tm6369)* and *set-2(bn129)* loss of function mutants. We observed that the level of H3K4me3 in *cfp-1(tm6369)* mutant is significantly reduced and was similar to reported *set-2(bn129)* allele (Figure 1B) suggesting that the *cfp-1(tm6369)* mutant is a loss of function allele.

**Figure 1.**
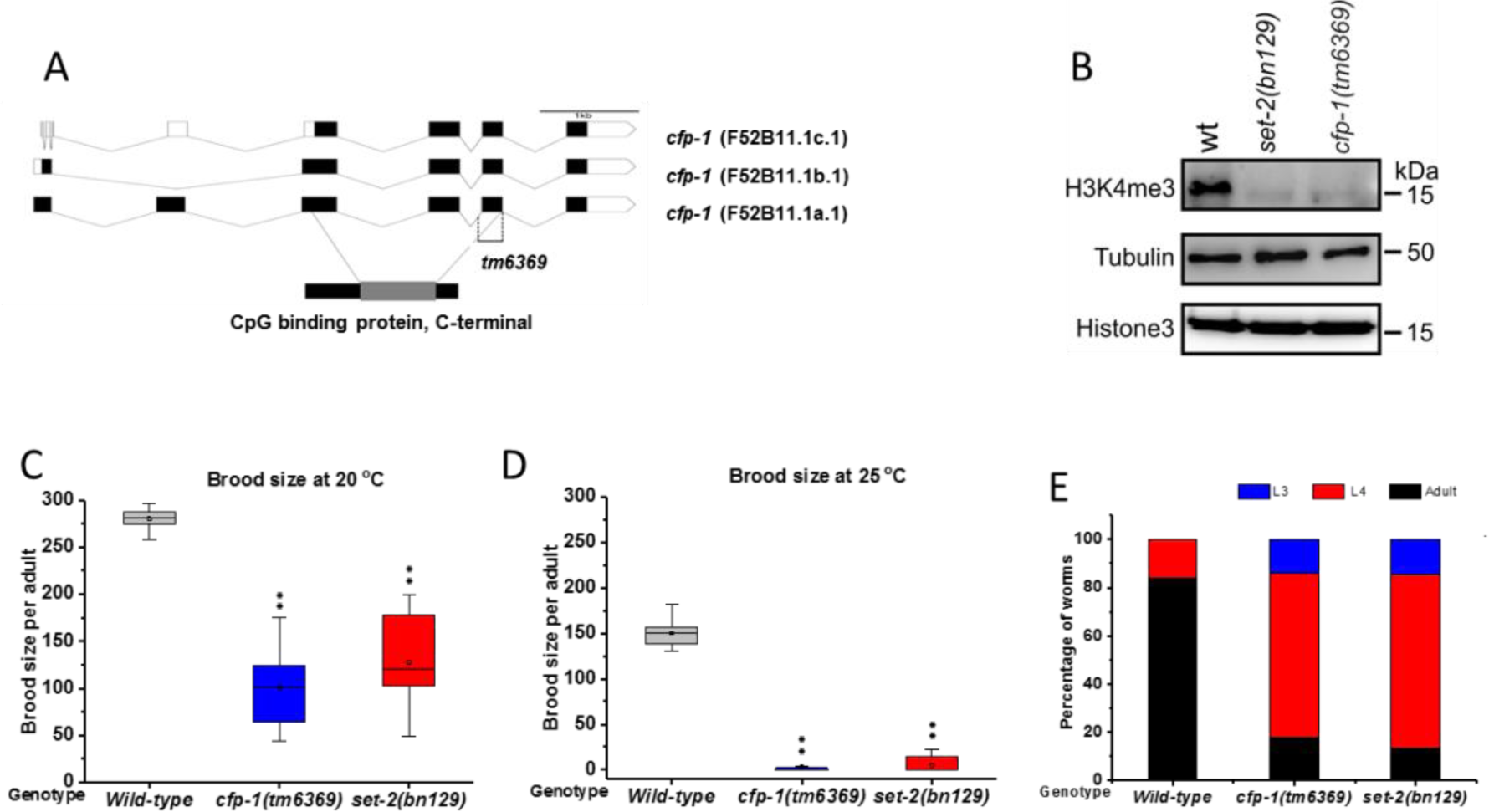
*cfp-1* (*tm6369*) is a loss of function allele. (A) Diagrammatic representation of *cfp-1(tm6369)* allele. 254bp were deleted from the exon 5 region of the F52B11.1a.1 transcript. The deleted region is indicated by dashed line. Black colour denotes the exon and grey colour denotes the CpG binding domain. (B) Western blot analysis, showing the reduced level of H3K4me3 in *cfp-1(tm6369)* and *set-2(bn129)* allele compared to wild type. Histone 3 (H3) and tubulin were used as a loading control. (C and D) Total brood size assay for wild-type *, cfp-1(tm6369)* and *set-2(bn129).*(C) *cfp-1(tm6369)* and *set-2(bn129*) average brood size was reduced compared to wild type at 20 °C. (D) Fertility was severely compromised at 25 °C, and 70% of *cfp-1(tm6369)* and *set-2(bn129)* mutants were sterile. (E) Developmental progress of *cfp-1(tm6369)*, *set-2(bn129)* and wild-type embryo monitored at 60 h at 20 °C. *cfp-1(tm6369)* and *set-2(bn129)* mutation displayed stochastic delays in development from embryo into young adults. P-values were calculated using student t-test: ** = P<0.01.

To explore the functional consequences of loss of *cfp-1* in *C. elegans* we carried the phenotypic characterisation of *cfp-1(tm6369)* mutant by conducting a fertility assay and measured the growth rate. For the fertility assay, we measured the brood size of *cfp-1(tm6369)* and *set-2(bn129)* mutants at two different temperatures. At 20 °C both mutants had a significant reduction in brood size compared to wild type and at 25 °C fertility was severely affected in both *cfp-1(tm6369)* and *set-2(bn129)* mutants (Figure 1C and 1D). In a previous study, it was reported that *set-2(bn129)* mutants display mortal germline phenotype, progressive loss of brood size over generations leading to sterility (Xiao et al., 2011). We also investigated the mortal germline phenotype of *cfp-1(tm6369)* mutant at 25 °C. Surprisingly, F2 generation of the *cfp-1(tm6369)* mutant at 25 °C was completely sterile. Taken together, these findings suggest that both *cfp-1* and *set-2* are equally important in maintaining the fertility.

We measured the growth rate of *set-2(bn129)* and *cfp-1(tm6369)* mutants and compared them to wild-type. *C. elegans* embryos pass through four larval development (L1, L2, L3 and L4) stages to reach adulthood. We measured the growth of freshly laid embryos of wild-type, *cfp-1(tm6369)* and *set-2(bn129)* mutants at 60 h. Both *cfp-1*(*tm6369*) and *set-2*(*bn129*) mutants show stochastic delays in development from embryo to adult (Figure 2E). After 60 h, ∼84% of wildtype embryos reached the adult stage, whereas only ∼18% of *cfp-1(tm6369)* and ∼18% of *set-2(bn129)* mutants were in the adult stage (Figure 1E). These results further strengthen that both *cfp-1* and *set-2* are required for the proper development of an organism.

**Figure 2.**
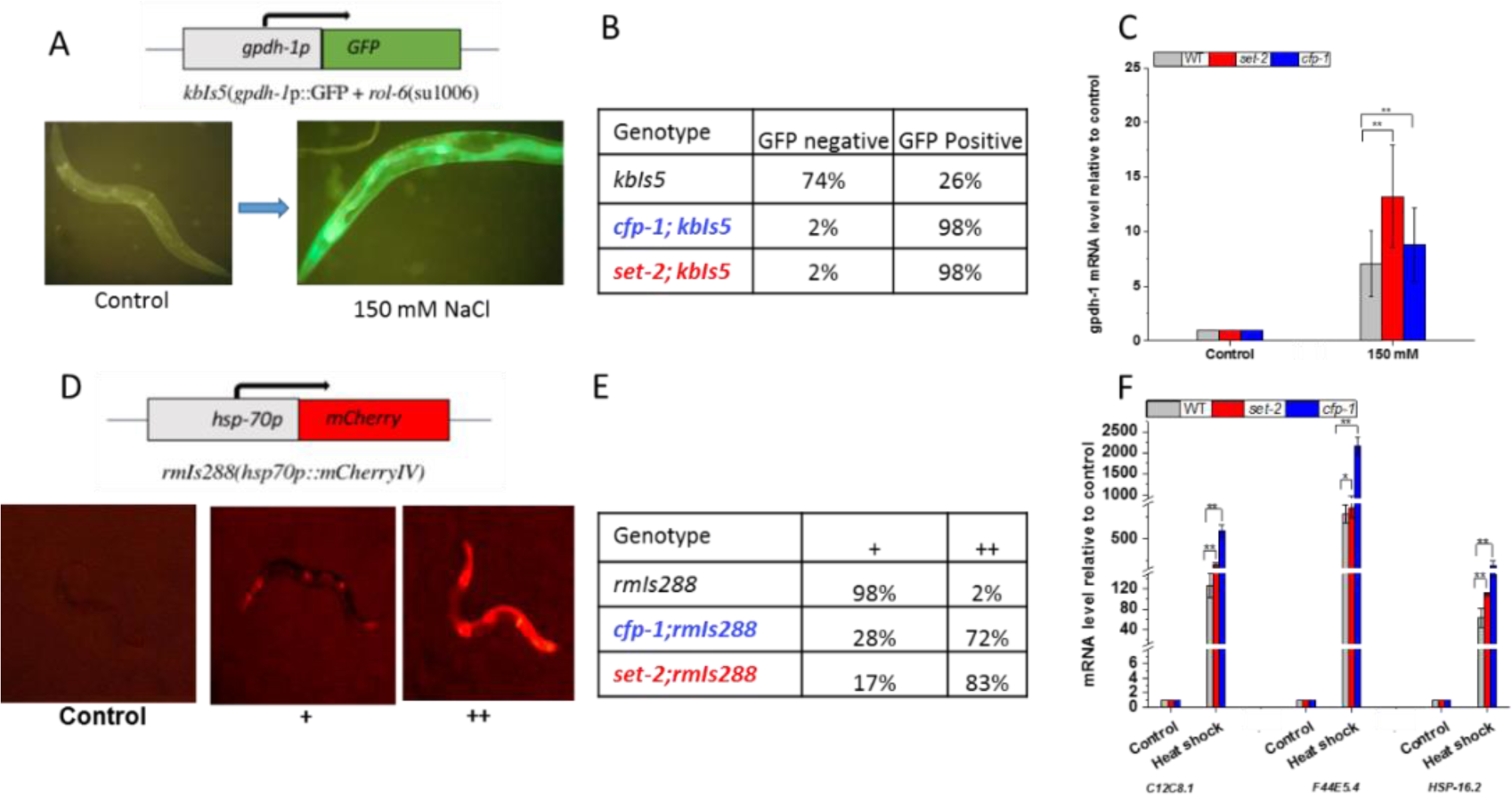
Loss of *cfp-1* and *set-2* results in stronger expression of inducible genes. (A) Vp198 (*gpdh-1p::GFP*) strain contains *GFP* downstream of *gpdh-1* promoter which is expressed in worms when shifted to a higher concentration of salt (150 mM). (B) Table showing the percentage of GFP positive and negative worms. L1 worms were grown at hypertonic condition (150 mM) for 72 h. COMPASS mutants show stronger GFP induction compared to *kbls5* (C) qPCR of *gpdh-1* gene at control (52 mM) and hypertonic condition (150 mM) in wild-type (grey), *cfp-1(tm6369)* (blue) and *set-2(bn129)* (red) mutants. *gpdh-1* expression level is higher in COMPASS mutants compared to wild-type. (D) AM722 (*hsp70p::mCherry*) strain contains mCherry downstream of *hsp-70* promoter which is expressed during heat shocks. (+) moderate expression, (++) stronger expression of mCherry. (E) Table showing the percentage of worms expressing mCherry. Worms were heat shocked at 35 °C for 1 h and left them to recover for 4 h. COMPASS mutants show stronger mCherry induction compared to *rmls288* (F) qPCR of heat shock genes *C12C8.1, F44E5.4* and *hsp-16.2* in wild-type (grey), *cfp-1(tm6369)* (blue) and *set-2(bn129)* (red) mutants before and after heat shock at 33°C for 1 h. *C12C8.1, F44E5.4* and *hsp-16.2* gene expression levels are higher in COMPASS mutants compared to wild-type. Figures are average of 2-3 biological replicates. P-values were calculated using student t-test: ** = P<0.01, * = P<0.05.

### *cfp-1* and *set-2* attenuate gene induction

We next investigated the role of *cfp-1* and *set-2* in gene expression by using reported salt-inducible reporter strain VP198 (kbIs5 [gpdh-1p::GFP + rol-6(su1006)]). VP198 contains green fluorescent protein (GFP) downstream of the *gpdh-1* gene promoter and is expressed in a higher salt environment (Figure 2A) (Lamitina et al., 2006). We crossed this strain with *cfp-1(tm6369)* and *set-2(bn129)* to generate *cfp-1(tm6369);kbIs5* and *set-2(bn129);kbIs5* double mutant strains and exposed them to a higher salt concentration (150 mM) than 52 mM which is required for growth. When exposed to hypertonic stress, both *cfp-1(tm6369);kbIs5* and *set-2(bn129);kbIs5* mutants displayed hyper-induction of the reporter gene compared to wild-type worms, as seen by higher GFP expression (Figure 2A and 2B). We also measured the endogenous expression level of *gpdh-1* gene at control levels (52 mM) and at higher salt (150 mM) concentrations in wild-type, *cfp-1(tm6369)* and *set-2(bn129)* mutants. We found that at higher salt concentration, the expression level of *gpdh-1* gene was 9 fold upregulated in *cfp-1(tm6369)* mutant and 13-fold in *set-2(bn129)* mutants compared to 7-fold in wildtype (Figure 2C).

To further investigate the role of *cfp-1* and *set-2* in gene induction regulation, we used heat shock reporter strain AM722 [*rmIs288(hsp70p::mCherry IV)].* AM722 contains mCherry downstream of heat shock promoter *hsp-70*, which is expressed during heat stress (Figure 2D) (Van Oosten-Hawle et al., 2013). We crossed this strain with *cfp-1(tm6369)* and *set-2(bn129)* to generate *cfp-1(tm6369);rmIs288* and *set-2(bn129);rmIs288* strains. We found that after heat shock mCherry expression was significantly higher in *cfp-1(tm6369);rmIs288* and *set-2(bn129);rmIs288* strains compared to *rmIs288* (Figure 2D and 3E). We also measured the endogenous expression of heat-inducible genes (*C12C8.1, F44E5.4* and *hsp-16.2*) in the *cfp-1*(tm6369) and *set-2(bn129)* mutant. *C12C8.1, F44E5.4* and *hsp-16.2* are heat inducible chaperones downstream of heat shock factor −1 (*hsf-1)* gene and are expressed during heat stress (Prahlad et al., 2008, Snutch et al., 1988, Jones et al., 1986). After heat shock at 33°C for one h, the expression of *C12C8.1, F44E5.4* and *hsp-16.2* were significantly upregulated in both *cfp-1(tm6369)* and *set-2(bn129)* mutants compared to wild-type (Figure 2F). Higher induction of heat and salt inducible genes in both *cfp-1(tm6369)* and *set-2(bn129)* mutants with a negligible level of H3K4me3 suggest that H3K4me3 can indeed play a repressive role in gene induction.

**Figure 3.**
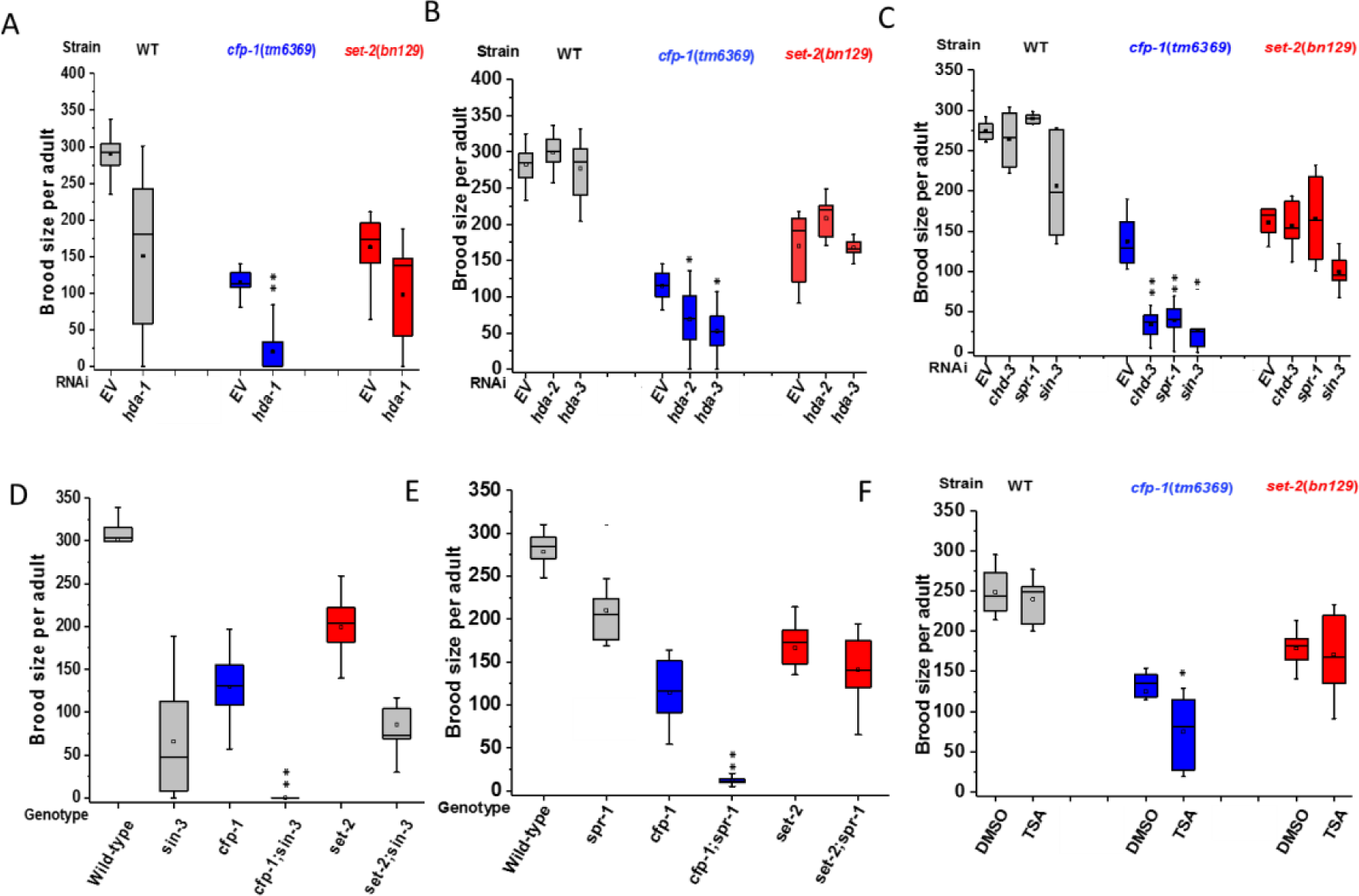
*cfp-1* genetically interacts with HDAC 1/2 to regulate fertility. Brood size assays showing genetic interactions between *cfp-1* and class 1/2 HDACs using RNAi knockdown (A-C), double mutants (D and E) and HDACs inhibitor (F). A multiplicative method was used to identify whether two genes interact to regulate fertility or not. For control, worms are fed on HT115 *E. coli* strain that empty RNAi feeding vector (EV). (A) Average brood size of *hda-1* RNAi. *hda-1* RNAi F1 resulted in higher percentage of embryonic lethality so fertility was assayed at P0. RNAi knockdown of *hda-1* resulted in a reduction in brood size in both wild-type and COMPASS mutants. Null hypothesis t-test (refer to the methods section) showed a synergistic interaction between *hda-1* and *cfp-1(tm6369)* but not *set-2(bn129).* (B and C) Average brood size on RNAi knockdown of *hda-2*, *hda-3, sin-3, spr-1* and *chd-3* in N2, *cfp-1*(*tm6369*) and *set-2(bn129)* mutants. Brood size of *cfp-1*(*tm6369*) was further reduced in these RNAi but had no significant impact on *set-2(bn129)* mutant brood size. (D) Average brood size of *sin-3(tm1276)* mutants. *cfp-1(tm6369);sin-3(tm1276)* double mutant was sterile. Brood size of *set-2(bn129);sin-3(tm1276)* was similar to the brood size of *sin-3(tm1276)* showing no genetic interaction. (E) Average brood size of *spr-1(ok2144)* mutants. Average brood size of *cfp-1(tm6369);spr-1(ok2144)* was lower than 10, whereas the brood size of *set-2(bn129);spr-1(ok2144)* was similar to *set-2(bn129).*(F) Brood size of wild-type, *cfp-1(tm6369)* and *set-2(bn129)* mutants treated with control (DMSO) or Trichostatin A (TSA). Average brood size of *cfp-1*(tm6369) mutant was slightly but significantly reduced when treated with TSA. Figures are average of 2-3 biological replicates. P-values were calculated using one-tailed student t-test: ** = P<0.01, * = P<0.05.

### CFP-1 genetically interacts with HDAC1/2 complexes

We conducted a mini RNAi screen to find the candidate genes that could either enhance or suppress the observed poor fertility phenotype of the *cfp-1(tm6369)* mutant. Previous studies have illustrated that crosstalk between COMPASS and histone acetylation plays an important role in ensuring proper gene regulation (Tang et al., Tie et al., 2009, Zhao et al., 2013), thus for the RNAi screen we selected histone acetyltransferases (*cbp-1, mys-4* and *hat-1*) and histone deacetylases (*hda-1, hda-2* and *hda-3)*. We conducted RNAi of selected HATs and HDACs on *cfp-1(tm6369)* and measured the effects on fertility. In the fertility screen, we did not observe any significant change in brood size of *cfp-1;mys-4* and *set-2;mys-4* double mutants, as well as RNAi knockdown of *hat-1* in COMPASS mutants (Supp. Figure 1A and B). *cbp-1* RNAi resulted in a larval arrest in WT, *cfp-1(tm6369)* and *set-2(bn129)* mutants so we could not measure the brood size. However, we did not observe significant changes in fertility of *cbp-1* gain of function mutant in *cfp-1* and *set-2* RNAi (Supp. Figure 1C). In contrast to HATs, RNAi knockdown of *hda-1, hda-2* and *hda-3* in *cfp-1(tm6369)* mutant significantly reduced the brood size, suggesting a synergistic genetic interaction between these histone deacetylases and *cfp-1* (Figure 3A and 3B). Interestingly, we found that unlike *cfp-1(tm6369), set-2(bn129)* brood size did not significantly reduce in RNAi of *hda-1, hda-2* and *hda-3* (Figure 3A and 3B).

We demonstrated that *set-2* and *cfp-1* play a similar role in fertility and development of *C. elegans*. However, we observed that RNAi knockdown of *hda-1, hda-2* and *hda-3* only enhance the low brood phenotype of *cfp-1(tm6369)* mutant. Different responses of the *cfp-1(tm6369)* and *set-2(bn129)* mutants to the same RNAi conditions could be due to differential sensitivity to RNAi. To investigate this, we carried out an RNAi sensitivity assay. We measured the RNAi sensitivity of *cfp-1*(tm6369) and *set-2(bn129)* mutants using *hmr-1, dpy-10* and *unc-15* genes with well-defined phenotypes. We found that both *cfp-1(tm6369)* and *set-2(bn129)* mutants responded similarly to the tested RNAi (Table 1). This suggests that the different response of *cfp-1(tm6369)* and *set-2(bn129)* mutants in *hda-1, hda-2* and *hda-3* RNAi background is not due to different sensitivity to RNAi.

**Table 1.** RNAi sensitivity assay: Sensitivity of wild-type, *set-2(bn129)* and *cfp-1(tm6369)* on RNAi was measured using *dpy-10*, *unc-15* and *hmr-1* RNAi. *dpy-10* was scored based on the average length of worms showing dumpy (shorter and fatter body morphology) phenotype. *unc-15* was scored based on the severity of uncoordinated phenotype (paralysis). *hmr-1* was scored based on the percentage of dead eggs of total brood.

| RNAi | Wild-type | *cfp-1(tm6369)* | *set-2(bn129)* |
| --- | --- | --- | --- |
| EV (Control) | - | - | - |
| *hmr-1* | <6% | <6% | <6% |
| *dpy10* | +++++ | +++++ | ++++ |
| *unc-15* | +++++ | ++++ | +++++ |

*hda-1, hda-2* and *hda-3* are the orthologs of mammalian class I HDACs (HDAC1/2) (Shi and Mello, 1998). HDAC1/2 are found in multiprotein complexes. Well characterised complexes containing HDAC1/2 are Sin-3, NuRD and CoREST, which contain *sin3*, *chd-3* and *spr-1* as a major subunit respectively (Solari and Ahringer, 2000, Choy et al., 2007, Bender et al., 2007, Gregoretti et al., 2004). To test which of these complexes interact with *cfp-1*, we carried out RNAi knockdown of *sin-3*, *chd-3* or *spr-1* in *cfp-1*(*tm6369*) mutant. Interestingly, RNAi knockdown of *sin-3*, *chd-3* or *spr-1*, further reduced the brood size of *cfp-1*(*tm6369*) mutant but there was no effect on the brood size of *set-2 (bn129)* (Figure 3C). To further confirm the RNAi results, we used *spr-1(ok2144)* and *sin-3(tm1276)* mutants. We found that all of the *cfp-1(tm6369);sin-3(tm1276)* double mutants were completely sterile and most of the *cfp-1(tm6369);spr-1(ok2144)* double mutants produced less than 10 progenies (Figure 3D and 3E). Similar to our RNAi results, we did not observe the synergistic interaction in *set-2(bn129);spr-1(ok2144) and set-2(bn129);sin-3(tm1276)* mutants. These findings support that *cfp-1* interacts with HDAC1/2 in a SET-2/COMPASS-independent manner.

We sought to investigate the functional link between *cfp-1* and HDAC1/2 complexes. One of the main functions of HDAC1/2 complexes is histone deacetylation, we asked if the inhibition of HDAC1/2 deacetylase enhance the low brood phenotype of *cfp-1(tm6369)* and *set-2(bn129)* mutant or not. We treated the *cfp-1(tm6369)* worms with Trichostatin A (TSA). TSA is a chemical that inhibits class I/II histone deacetylase and TSA treated cells have a significant gain in histone acetylation (Crump et al., 2011). TSA is a toxic chemical, thus we used 4 mM which is an established non-toxic dose for *C. elegans* (Vastenhouw et al., 2006). We found that the average brood size of wild-type and *set-2(bn129)* mutant were not affected. In contrast, the brood size of *cfp-1(tm6369)* mutant was slightly but significantly reduced (Figure 3F). This further confirms the genetic interaction between *cfp-1* and HDAC1/2 and provide the functional link between *cfp-1* and HDACs.

### *cfp-1* and *set-2* independently regulates fertility and growth

We did not observe any genetic interaction between *set-2* and tested HDACs, but we found a clear genetic interaction between *cfp-1* and HDAC1/2 complexes. This finding suggested that *cfp-1* and *set-2* might act in separate pathways to regulate fertility. To investigate this, we generated *cfp-1*(*tm6369);set-2(bn129)* double mutant and measured the brood size. If both *cfp-1* and *set-2* were acting in a similar pathway then the average brood size of double mutants should be similar to single mutants. Interestingly, we found that the average brood size of *cfp-1*(*tm6369);set-2(bn129)* double mutant was significantly lower than the average brood size of *cfp-1*(*tm6369)* and *set-2(bn129)* single mutants (Figure 4A). This clearly suggests that *cfp-1* and *set-2* act in separate pathways to regulate fertility.

**Figure 4.**
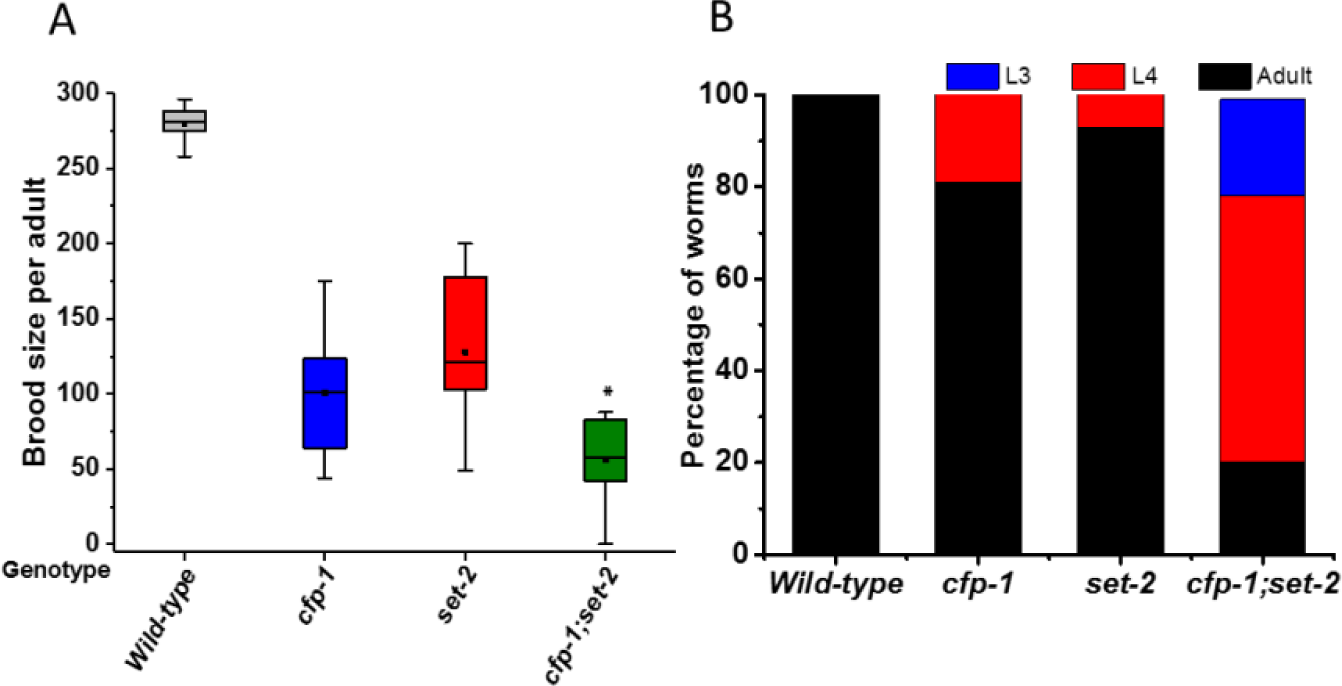
*cfp-1* and *set-2* act in a different pathways. (**A**) Brood size of wild-type, *cfp-1(tm6369), set-2(bn129)* and *cfp-1(tm6369);set-2(bn129)* mutants at 20 °C. Double mutants have significantly reduced brood size compared to single mutants, however the difference in brood size is not synergistic (based on Null hypothesis T-test). (B) Developmental progress of wild-type, *cfp-1(tm6369), set-2(bn129)* and *cfp-1(tm6369);set-2(bn129)* embryo monitored at 68 h at 20 °C. *cfp-1(tm6369);set-2(bn129)* grow slower than *cfp-1(tm6369)* and *set-2(bn129)* single mutants. Figures are average of two biological replicates. P-values were calculated using one-tailed student t-test: * = P<0.05.

We also carried out growth kinetics of *cfp-1*(*tm6369);set-2(bn129)* double mutant and compared to *cfp-1*(*tm6369)* and *set-2(bn129)* single mutants. We found that the double mutants grow slower compared to single mutants (Figure 4B). Taken together, these findings suggest that even though *cfp-1* and *set-2* are key subunits of COMPASS complex, they also act in separate pathways in *C. elegans* development.

## Discussion

Over the past decade, various research groups have emphasized the importance of CFP-1 in cell fate specification and cell differentiation. However, the contribution of CFP1 to gene regulation is not fully understood. In this study, we set out to elucidate the impact of the loss of CFP-1 on gene induction and development by using *cfp-1(tm6369)* and *set-2(bn129) C. elegans* mutants. Phenotypic characterisation of *cfp-1(tm6369)* and *set-2(bn129)* mutants suggests that *cfp-1* and *set-2* play an important role in fertility and development of *C. elegans*. We found that in *cfp-1(tm6369)* and *set-2(bn129)* mutants, the induction of heat and salt inducible genes were significantly higher than the wildtype. The similar function of *cfp-1(tm6369)* and *set-2(bn129)* in fertility and in gene induction supports that CFP-1 function in a COMPASS complex. However, we also found that *cfp-1* and *set-2* act in separate pathways to regulate fertility, and that *cfp-1* but not *set-2* genetically interacts with HDAC1/2 complexes to regulate fertility. These findings suggest a function of *cfp-1* outside of the Set1/COMPASS complex. We propose that CFP-1 could interact with COMPASS complex and HDAC1/2 complexes in a context dependent manner (Figure 5).

**Figure 5.**
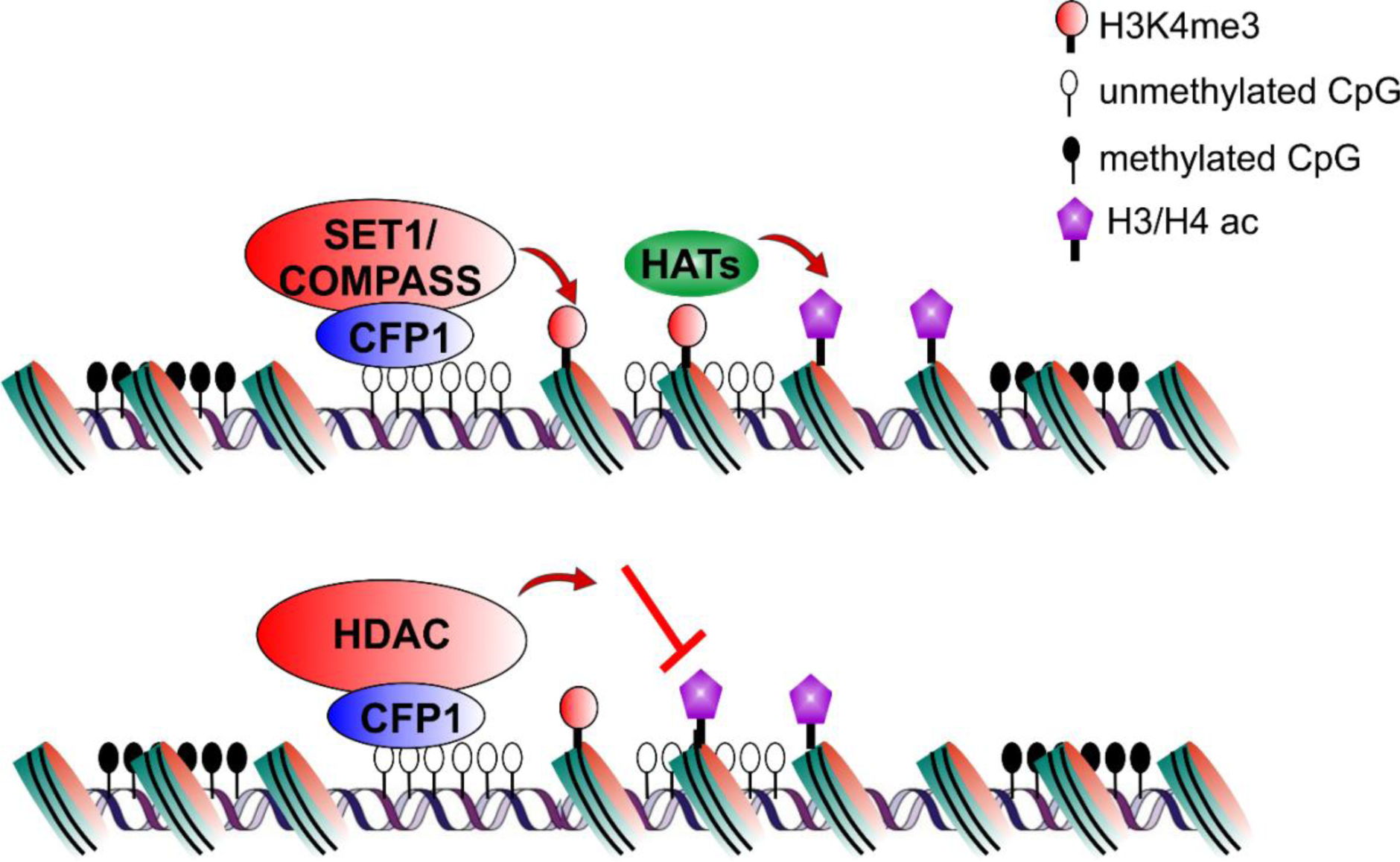
Proposed model indicating that CFP-1 cooperates with SET-2/COMPASS and/or with HDACs in a context dependent manner. Canonical function of CFP-1 is to recruit SET-2/COMPASS complex at promoter regions by binding into unmethylated CpG island. Non-canonical function of CFP-1: CFP-1 could also recruit HDAC complexes at promoter region to deacetylate the histones. Based on the physiological condition CFP-1 could either interact with the COMPASS complex or with HDAC complexes.

CFP-1 and SET-2 are major subunits of COMPASS complex responsible for bulk H3K4me3 (Simonet et al., 2007, Li and Kelly, 2011, Xiao et al., 2011). Here we observed that loss of *cfp-1* or *set-2* results in a dramatic reduction of the H3K4me3 level. We also observed the hyper induction of salt and heat inducible genes following the loss of *cfp-1* or *set-2.* The observed hyperinduction could be due to an increase in chromatin accessibility in the loss of H3K4me3. This could be supported by the fact that in yeast H3K4me2/3 repress *GAL1* gene induction by recruiting histone deacetylase complex called RPD3S (Margaritis et al., 2012). Similarly, another study suggests that H3K4me3 acts as a memory to repress the *GAL1* reactivation by recruiting Isw1 ATPase which limits the RNA polymerase II activity (Zhou and Zhou, 2011). It could be possible that the H3K4me3 could interact with repressor complex to restrict the binding of transcription factors such as HSF-1. Alternatively, H3K4me3 could regulate gene induction by altering the regulation of RNA polymerase II (Pol II) pausing. The previous study in *Drosophila melanogaster* has shown that paused Pol II is found in the promoter of *hsp* genes of un-induced drosophila and are primed for transcription activation in response to a stimulus (O’brien and Lis, 1991). It is possible that H3K4me3 at promoters could act as a regulator to maintain paused Pol II and prevent the burst of transcription before induction.

Another key finding of this study is that *cfp-1* genetically interacts with HDAC1/2 complexes. We found that RNAi knockdown of key subunits (*hda-1*, *hda-2*, *hda-3*, *chd-3*, *sin-3* and *spr-1*) of HDAC1/2 complexes enhanced the low brood size phenotype of *cfp-1(tm6369)* mutant. Surprisingly, we did not observe any genetic interaction between HDAC1/2 complexes and *set-2* suggesting the Set1/COMPASS independent function of CFP-1. Furthermore, we found that a decrease in deacetylase activity enhances the low brood phenotype of *cfp-1(tm6369)* mutant suggesting that CFP-1 might cooperate with HDACs for deacetylase activity. Previous study in yeast has shown that *Spp1* (yeast ortholog of *cfp-1*) exist in Mer2-Spp1 complex (Adam et al., 2018). It could be possible that CFP-1 is also present in HDAC complexes. As shown in Figure 5, CFP-1 could interact with both Set1/COMPASS and/or HDACs complexes in a context dependent manner.

## Materials and method

### Strains and their Maintenance

The following strains were used for experimental purpose. N2(wild-type), *set-2(bn129), cfp-1(tm6369), mys-4(tm3161), set-2(bn129);mys-4(tm3161), cfp-1(tm6369);mys-4(tm3161), cbp-1(ku258), rmIs288, cfp-1(tm6369);rmIs288*, *set-2(bn129);rmIs288, kbIs5[gpdh-1p::* GFP *+rol-6(su1006)], cfp-1(tm6369);kbIs5, cfp-1(tm6369);kbIs5, spr-1(ok2144), set-2(bn129);spr-1(ok2144), cfp-1(tm6369);spr-1(ok2144), sin-3(tm1276), set-2(bn129);sin-3(tm1276)*, and *cfp-1(tm6369);sin-3(tm1276).* Worms were maintained at 20 °C unless stated at standard growth condition. They were grown on *Escherichia coli* OP50 seeded Nematode Growth Medium (NGM) petri plates.

### Western Blot

Starved L1 (3.2-3.5K) worms were pelleted in M9 and snap-frozen at −80 °C. Pellets were recovered in lysis buffer (50 mM Tris-Cl (pH 8), 300 mM NaCl, 1 mM PMSF, 1 mM EDTA, 0.5% Triton X-100 and protease inhibitor cocktail (Xiao et al., 2011) and sonicated at 20% amplitude. Protein concentration was measured by using the Bradford method. 50 µg of total protein was loaded in each well. The protein was transferred to the nitrocellulose membrane. The membrane was cut into two part based on the molecular weight of tubulin ∼50 kDa, and H3 ∼15 kDa. Membranes were incubated with 1:5000; anti-H3K4me3, 1:5000; anti-H3 or 1:5000; anti-tubulin antibodies. Since H3 and H3K4me3 were of similar molecular weight, first H3K4me3 was detected followed by stripping the membrane for H3 detection. The membrane was washed twice and incubated with 1:5,000 dilutions of HRP-linked secondary antibodies. H3 and tubulin were used as a loading controls.

### Brood size assay

For brood size at 20 °C, ten L4 worms were picked and transferred to individual plates (1 worm per plate). Worms were transferred into new plates every day until laying ceased. Old plates were counted for a total number of eggs and were stored at 20 °C for two consecutive days and subsequently, scored for the number of live progeny. Animals that crawl out of the plates and lost were not included (Xiao et al., 2011). This experiment was repeated three times.

For Brood size at 25 °C, twenty L4 worms were picked from 20 °C and transferred to new OP50 seeded plates. They were allowed to lay eggs for overnight at 25 °C. Next day, all mother worms were picked and transferred to new plates and left for 5-6 h. Mothers from new plates were removed, and eggs were allowed to reach L4 at 25 °C. From the new plate, ten L4 worms were picked and transferred to an individual plate. Worms were transferred into new plates every day until they stop laying. Old plates were counted for a total number of eggs on plates and were stored at 25 °C for two consecutive days and subsequently scored for the number of live progeny. Animals that crawled out of the plates and lost were not included (Xiao et al., 2011).

Student T-test was performed to investigate the potential interaction between two genes. Under null hypothesis, where no genetic interaction between two genes is assumed, the expected brood size of the double mutants (or RNAi knockdown of a gene in a single mutant) is the product of the brood size of the single mutants (or single mutant and the RNAi knockdown of the gene in a wild-type) divided by the average brood size of wildtype. A one-sided T-test is done to compare the expected (under null-hypothesis) brood size with the observed brood size of double mutants (or RNAi knockdown of a gene in a single mutant).

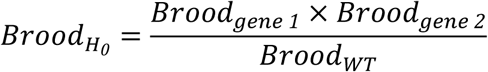

*Brood_H0_* = Expected Brood size of double mutant (or RNAi) under the null hypothesis

*Brood_gene1_* = Actual Brood size of first mutant (or RNAi)

*Brood_gene2_* = Actual Brood size of second mutant (or RNAi)

*Brood_WT_* = Actual Brood size of Wild-type

### RNAi sensitivity assay

Three L3/L4 worms are transferred from OP50 plates to EV, *dpy-10, unc-15* and *hmr-1*. RNAi plates. Worms are left to grow for 48 hours before being transferred to fresh RNAi plates. After 24 hours the worms are transferred again to a fresh RNAi plate. Brood size is counted for each plate and the sum is divided by three to give average brood size of the worm as a control. Severity of phenotype in *dpy-10* RNAi was assessed by comparing the body length of mutant worms with N2 in *dpy-10* RNAi. For *unc-15*, number of adult worms that are able to move their body are counted. For *hmr-1*, percentage of dead eggs are observed.

### Growth Kinetics assay

Twenty synchronised L4 worms were picked from 20 °C and transferred to new OP50 seeded plates. They were allowed to lay eggs for overnight at 20 °C. Next day, all mother worms were picked and transferred to new plates to lay eggs and left for 5-6 h. Mothers from the new plates were removed, and eggs were left to grow for 60 or 68 h. After the respective time, worms were transferred to the tubes, washed twice with M9, frozen in methanol for 1 h, DAPI stained and observed under normaski.

### Heat shock experiment

For reporter assay, synchronized first-day young adult worms grown at 20 °C were heat shocked at 35 °C for 1 h and left them to recover for 4 h. Worms were observed using an RFP filter on a Leica MZ10 F fluorescence microscope for the expression of mCherry. For qPCR, synchronized first-day young adult worms grown at 20 °C were heat shocked at 33 °C for 1 h. After heat shock worms were collected, washed twice with M9 and snap frozen at −80 °C.

### Salt induction experiment

For reporter assay, starved L1-stage worms were placed on NGM plates containing 52 mM and 150 mM NaCl. After 72 h worms were observed under a fluorescence microscope for the expression of GFP. For qPCR, starved L1-stage worms were placed on NGM plates containing 52 mM and 150 mM NaCl. After 72 h worms were collected, washed twice with M9 and snap frozen at −80 °C.

### RNAi Screening

Indicated RNAi clones were streaked on plates containing ampicillin (100 μg/mL) and tetracycline (100 μg/mL) and incubated overnight at 37 °C. The overnight culture was inoculated in a 2ml LB with ampicillin (100 μg/mL) and incubated for 6-8 h at 37 °C in a shaking incubator. The grown bacterial culture was seeded on a dried NGM plate containing 1mM IPTG and 100 μg/mL ampicillin. Seeded plates were dried at room temperature then incubated for 24 h at 37 °C. To all RNAi experiment except for *hda-1*, L1 worms were spotted on RNAi plates, and their progeny(F1) were used for the experiments. For *hda-1* RNAi, spotted L1 (P0) were used for all the experiments.

### Fertility assay of TSA treated worms

NGM plates containing 4mM Trichostatin A (SIGMA) or Dimethyl sulfoxide (DMSO) were prepared. L1(P0) worms were transferred into TSA or DMSO plates and incubated at 20 °C. Ten L4 worms (F1) were picked and transferred to an individual TSA or DMSO plate. and their fertility was assayed.

### RNA extraction and qPCR

RNA was extracted using Direct-zol RNA miniprep. Extracted RNA was reverse transcribed to obtain cDNA using iScript cDNA synthesis kit (Bio-Rad). qPCR was performed with SYBR^®^ Green (Biorad). Fold change in (*C12C8.1, F44E5.4* and *hsp-16.2*) heat shock genes and *gpdh-1*(salt inducible gene) was measured using 2^−ΔΔCt^ formula. *tba-1* and *pmp-3* were used as a reference gene to calculate the fold change. qRT-PCR was performed on two biological replicates.

## Acknowledgements

We thank Dr. Ron Chen for his support and advice on the project. We thank James Cain for the technical support. We thank Dr Patricija van Oosten-Hawle for providing the AM722 strain. We thank Dr. Amit Kumar, Laura Warwick and Dovile Milonaityte for careful reading of the manuscript.

## Author’s contribution

B.P. designed and performed the majority of experiments. Y.C. performed RNAi sensitivity assay and J.B. performed heat shock reporter assay. B.P. prepared the manuscript and all authors read and approved the final manuscript.

## Supplementary figures

**Supplemental Figure 1.**
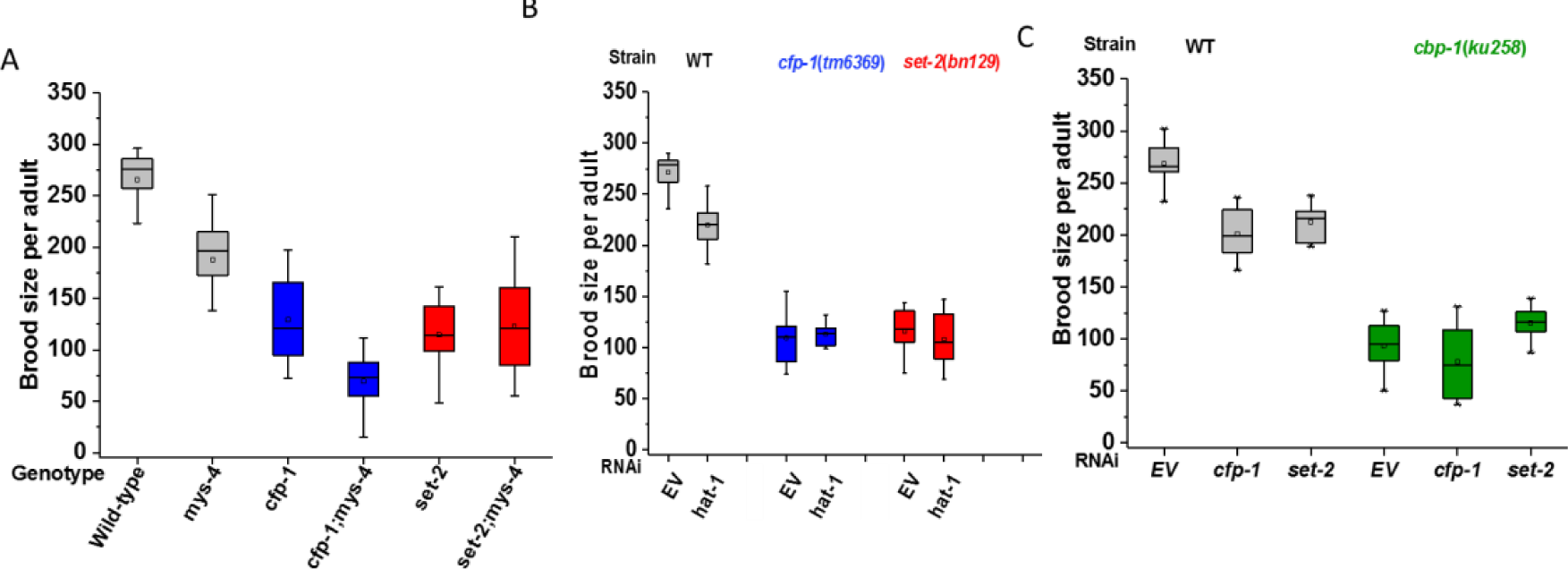
*cfp-1* and *set-2* do not interacts with HATs to regulate fertility. (A) Brood size of wild-type, *cfp-1(tm6369)*, *set-2(bn129)*, *cfp-1(tm6369);mys-4(tm3161)* and *set-2(bn129);mys-4(tm3161)* mutants at 20 °C. *cfp-1(tm6369);mys-4(tm3161)* double mutant had reduced brood size compared to single mutants, however the difference in brood size is not synergistic (based on Null hypothesis T-test)..(B) Brood size of wild-type, *cfp-1(tm6369)* and *set-2(bn129)* mutants on *hat-1* RNAi. RNAi knockdown of *hat-1* did not had significant impact on the brood size of *cfp-1(tm6369)* or *set-2(bn129)* mutants. (C) Brood size of *cbp-1(ku258)* on RNAi of EV, *cfp-1* and *set-2*. Brood size of *cbp-1(ku258)* on RNAi of *cfp-1* and *set-2* was similar to EV. Figures are average of two-three biological replicates.

